# Borrowed real estate: *Sappinia lukoli*, a new species of dung-dwelling amoeba that aggregates and hijacks the fruiting bodies of phylogenetically distant sorocarpic protists

**DOI:** 10.64898/2026.08.10.743969

**Authors:** Tristan C. Henderson, Benjamin Mixon, Charles Thompson, Cassandra F. van Riessen, Matthew W. Brown

## Abstract

Upon defecation, dung enters the world as a short-lived bounty of nutrients. Yet, it becomes increasingly hostile as it ages. In two days dung can be dominated by predatory insect larvae, mites, nematodes, zoopagalean fungi, and toxin-producing bacteria. With rapidly changing chemical composition and dehydration, this environment becomes inhospitable to the life it originally hosted. It is in these contexts that we see a remarkable pattern in dung’s protist diversity: across at least four eukaryotic supergroups, dung-dwelling amoeboid species have independently evolved cooperative behaviors by which cells navigate to the surface and form multicellular aggregates. Here we present a nuanced case of this behavioral diversity by describing *Sappinia lukoli*, a new amoeba species within Amoebozoa isolated from cattle dung. Other *Sappinia* species tend to be large and able to ‘stand’ by pushing their cell bodies into the open air. *S. lukoli* is the smallest *Sappinia* species described to date and does not stand. Instead, its cells aggregate at the distal tips of dung fibers and remain there as the culture ages. We also find that *S. lukoli* eats other dung-dwelling protists such as *Sorodiplophrys stercorea* (supergroup Stramenopiles) and *Guttulinopsis vulgaris* (supergroup Rhizaria). Strikingly, *S. lukoli* will gather inside the multicellular fruiting bodies built by *S. stercorea* and *G. vulgaris* on the dung surface. The cells of *S. lukoli* pack between host spores, effectively hijacking their fruiting bodies and gaining access to dispersal vectors. To our knowledge, this is the first record of a protist colonizing the aggregative fruiting bodies of other protists across multiple eukaryotic supergroups. *S. lukoli*’s own aggregation is yet another independent origin of this behavior in dung, and we propose that the habitat itself repeatedly selects for cooperation among its microbial residents.

## 1. Introduction

The genus *Sappinia* (Amoebozoa: Discosea: Thecamoebida) is comprised of free-living naked amoebae that are well characterized by their two closely appressed nuclei and membranous wrinkles running down the length of their cells. Coating their cell membrane is a dense layer of alcianophilic sugars, which may help protect against desiccation (Goodfellow et al., 1974). This layer may matter because most *Sappinia* strains live in terrestrial microhabitats that dry out, such as bark, moss, soil, dead-plant matter, and the dung of many animals such as cows, horses, bison, bats, and penguins (Noble, 1958; Mulec et al., 2016; Tyml and Dyková, 2018). Four species are currently named, *S. pedata, S. diploidea, S. platani*, and *S. dangeardi*, alongside multiple molecularly distinct, undescribed lineages (Wylezich et al., 2009, 2015; Henderson et al., 2024). Cases of human amoebic encephalitis have been associated with the genus, first reported as *S. diploidea* (Gelman et al., 2001) and later molecularly re-identified as *S. pedata* (Qvarnstrom et al., 2009), with a second, as-yet-unconfirmed *S. pedata* case reported from Kerala, India in 2025 (Gowdham, 2025). However, most *Sappinia* isolates to date have been from natural settings and not seen as parasitic. The most recent *Sappinia* phylogeny shows several branches of undescribed diversity, specifically near *S. diploidea*, representing undescribed lineages from soil and soil-adjacent environments like dung (Henderson et al., 2024). From this and other literature, we propose that animal dung continues to harbor undescribed microbial diversity, behaviors, and ecological interactions (Noble, 1958; Bass et al., 2016; Tice et al., 2016).

Protists in dung encounter many selective pressures that favor escape strategies. Dung is ephemeral, rapidly desiccating, frequently visited by arthropods (Sladecek et al., 2013), and patrolled by sub-surface zoopagalean fungi that prey selectively on protists (Drechsler, 1939; Blackwell and Malloch, 1991). Protists in terrestrial environments, especially dung, answer these conditions with a conspicuous tendency to elevate themselves above their substrate. Most often this is done by making spore-bearing fruiting bodies, constructed in different ways by protosteloid amoebae, myxomycetes, and cellular slime molds (Shadwick et al., 2009; Fiore-Donno et al., 2010). An alternative to fruiting bodies is seen in terrestrial *Sappinia*, which often display ‘standing’ behavior. Here, cells navigate to the surface of their substrate, anchor on it, and form an actin-filled ‘leg’ that actively holds most of their cell mass in the open air (Henderson et al., 2024). Together, all these behaviors are thought to aid in dispersal. Standing cells have been shown to attach firmly to phoretic gamasid mites and beetles that travel between dung patches (Blackwell and Malloch, 1991). Insect vectors, for example, can also pick up fruiting bodies or standing cells and carry them to more favorable conditions, as has been shown with *Dictyostelium discoideum* (Smith et al., 2014).

The selective pressures posed by dung (i.e. predation, desiccation, and chemical stress) may also favor cooperation (Tong et al., 2022). Dung environments are unusually rich in microbial eukaryotes that have independently evolved aggregative behaviors. Such is the case with sorocarpic amoebae, whose cells coordinate in the hundreds to thousands to construct macroscopic spore-bearing fruiting bodies. This behavior has independently evolved at least seven times across the eukaryotic tree (Brown and Silberman, 2013; LamŻa, 2023), with the majority of lineages being from dung, including *Copromyxa protea* (Amoebozoa; Brown et al., 2011), *Sorodiplophrys stercorea* (Stramenopiles; Tice et al., 2016), *Guttulinopsis vulgaris* (Rhizaria; Brown et al., 2012), *Fonticula alba* (Holozoa; Brown et al., 2009), and various dictyostelids (Amoebozoa; Brefeld, 1869; van Tieghem, 1880). This suggests a strong selection for cellular cooperation in this habitat. Species within *Sappinia* may be a part of this greater pattern. Certain dung-dwelling *Sappinia* have been recorded to aggregate, forming clusters or ‘heaps’ of dormant cells at the distal tips of small substrate projections (Cienkowski, 1873; Dangeard, 1896; Olive, 1902; Cook, 1939; Raper, 1960; Brown et al., 2007; Henderson et al., 2024). However, aggregation in *Sappinia* is not well described and is not considered sorocarpy. More broadly, how these dung-dwelling protists cooperate, discriminate between genetic kin, and interact with each other has yet to be investigated.

Here we describe *Sappinia lukoli* sp. nov. isolated from cattle dung at Mississippi State University. *S. lukoli* is distinct in phylogenetic position, morphology, and behavior. Cells of *S. lukoli* are 2-3 times smaller in length than other *Sappinia* species and do not stand like other *Sappinia* from dung. Instead, its cells aggregate at the distal ends of dung fibers and remain so as the culture ages. We show that *S. lukoli* preys on other dung-dwelling amoebae, such as *Sorodiplophrys stercorea* and *Guttulinopsis vulgaris*. Strikingly, *S. lukoli* will preferentially gather within the fruiting bodies of these two sorocarpic amoebae, between the host spores, effectively hitching a ride if picked up by a vector. This generalist “hijacking” of sorocarpic protist fruiting bodies across at least two eukaryotic supergroups represents a suite of new cryptic interactions to explore between the microbial inhabitants of dung environments.

## 2. Results

### 2.1 Discovery, morphology, and phylogenetic placement of *Sappinia lukoli* sp. nov

On the surface of our cow dung samples, we picked a single sorocarp of *Sorodiplophrys stercorea* (strain SsSF25) and transferred it to sterilized dung. Scanning the culture showed that the sorocarps of this *S. stercorea* had an abnormal studded appearance. Upon closer inspection, we found hundreds of *Sappinia* cells dispersed within these sorocarps, characterized by their paired nuclei. After, we established a clonal strain of this *Sappinia* from a single cell, maintained it in co-culture with *S. stercorea*, and successfully froze the culture in liquid nitrogen storage. In the following paragraphs, we show through morphological and phylogenetic characterization that this *Sappinia* isolate represents a previously undescribed species that displays aggregative behavior. Considering that this strain was isolated from the homelands of the Choctaw Native American people, we use the Choctaw word “lukoli” meaning “clustered,” “grouped,” or “gathered” to herein name this species *Sappinia lukoli* sp. nov.

*Sappinia lukoli* cells have the characteristic *Sappinia* features; a monopodial lingulate ovate shape, lateral folds, and paired nuclei (Fig. 1). They also form multiple small contractile vacuoles that merge before expelling their contents, as has been observed in *S. dangeardi*. However, *S. lukoli* is the smallest *Sappinia* species described to date, both in cell size and nuclear diameter (Table 1). Their cells often have dorsal longitudinal wrinkles during movement (Fig. 1F), which is a more describe characteristic of their sister genus *Thecamoeba*. Additionally, *S. lukoli* cells do not stand when given the same conditions that trigger *S. dangeardi* and *S. pedata* to stand on dung. Including its distinct morphology, this strain is distinguished from other *Sappinia* by its sequence divergence and phylogenetic position. We sequenced the full-length nuclear encoded small subunit rRNA gene (SSU) of *Sappinia lukoli* strain SappSF25 (2,525 bp). Uncorrected pairwise distances (Suppl. Table 1) show that it differs from all full-length *Sappinia* SSU sequences by at least 5.4% (≤ 94.6% identity), the closest being *Sappinia diploidea* strain Vb (EU881939). In the genus, *S. lukoli* groups with *S. diploidea, S. dangeardi*, and several undescribed strains (Fig. 2). This topology is consistent with other published 18S analyses of the genus (Henderson et al., 2024; Corsaro et al., 2017).

**Table 1.** Measurements of motile cells across *Sappinia* strains. All dimensions in µm. Ranges are observed from min to max, not as mean-derived intervals. Nucleus is the diameter of a single nucleus. n = cells measured. n.r. = not reported. Morphometrics and each source’s original wording are in Supplementary Data 1.

| Species | Strain | Length | Breadth | Nucleus | n | Source |
| --- | --- | --- | --- | --- | --- | --- |
| *S. lukoli* | SappSF25 | 18.8–28.2 | 11.4–18.6 | 2.29–3.51 | 75 | this study |
| *S. pedata* | MSU2206 | 42–91 | 23–54 | n.r. | 129 | Henderson et al., 2024 |
| *S. dangeardi* | BF22-2A | 47–117 | 26–63 | 4.5–7 | 161 | Henderson et al., 2024 |
| *Sappinia* sp. | SG10G | 46.1–56.4 | 20.5–35.8 | n.r. | 20 | Tymł and Dyková, 2018 |
| *S. platani* | PL-247 (CCAP 1575/4) | 57–76 | 23–38 | ca 4 | 10 | Wylezich et al., 2015 |
| *S. pedata* | UK05-34-3aM | 35–85 | 16–42 | ca 4.5 | 20 | Brown et al., 2007 |
| *S. diploidea* | Sd-plat-4/02 (CCAP 1575/2, neotype) | 55–60 | n.r. | 3.8–4.6 | n.r. | Michel et al., 2006 |
| *S. diploidea* | CCAP 1575/1 | 43–86 | n.r. | 5.9–6.9 | 100 | Goodfellow et al., 1974 |

**Fig. 1.**
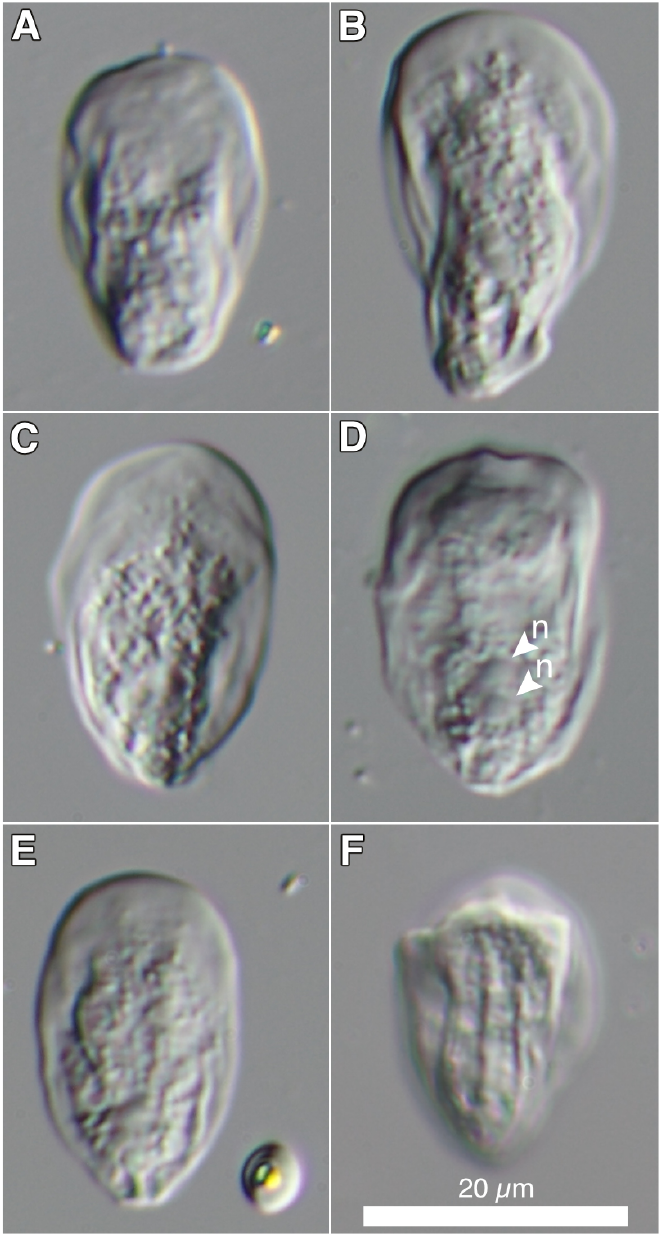
Microscopy of *Sappinia lukoli* sp. nov. (A–F) Locomotive trophozoites moving on a glass slide, showing the monopodial, lingulate-to-ovate shape and lateral folds characteristic of the genus. (D) Arrowheads (n) mark the two closely appressed nuclei. (E) A single *Sorodiplophrys stercorea* spore is also visible (lower right). (F) Dorsal folds of a moving cell. All images differential interference contrast (DIC). Scale bar = 20 µm.

**Fig. 2.**
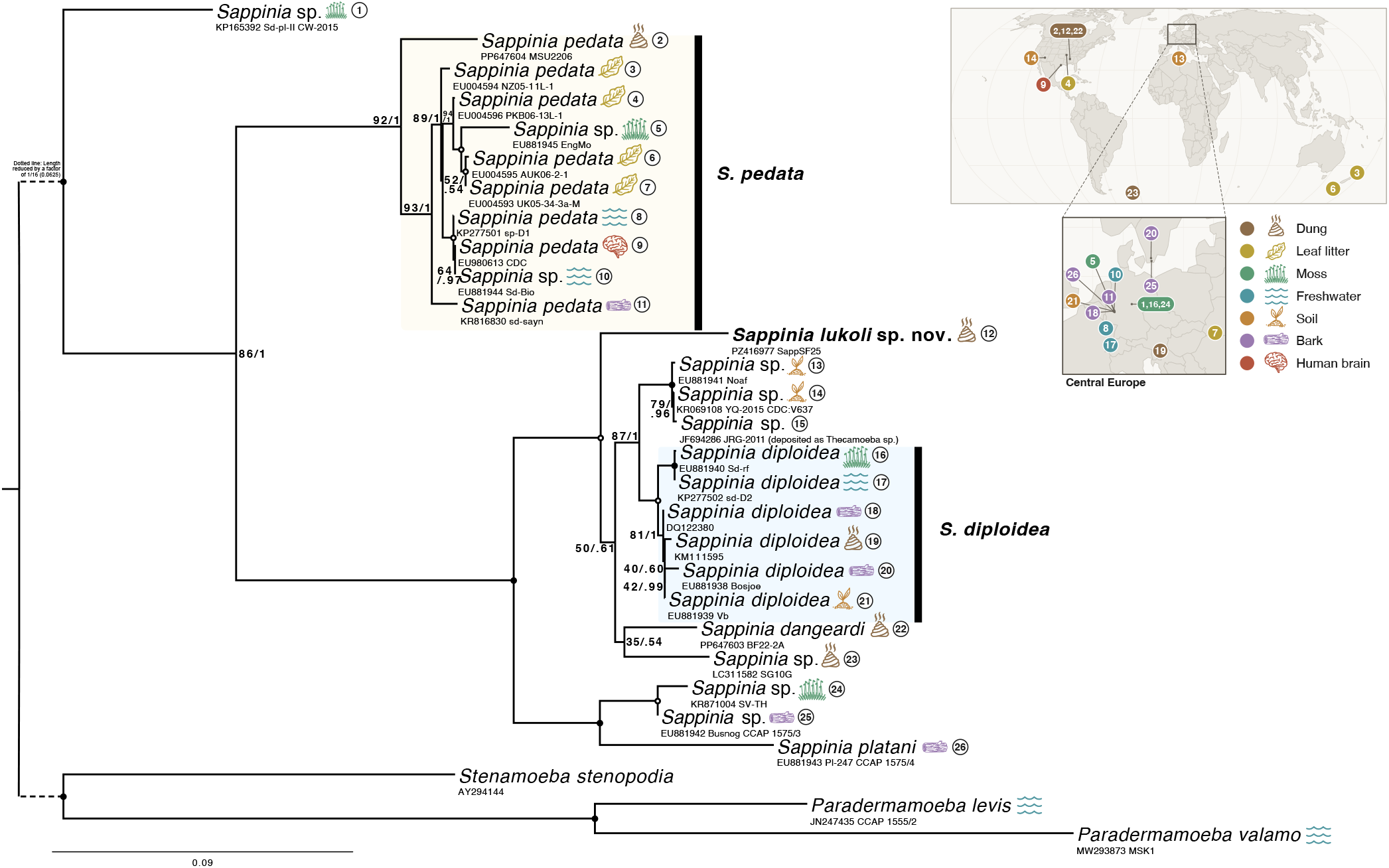
Maximum likelihood SSU (18S) rRNA gene tree of the genus *Sappinia* (Amoebozoa, Discosea, Thecamoebida). The tree was inferred in RAxML v8.2.12 under the GTR + Γ + I model from 2,368 aligned nucleotide positions with 1,000 rapid bootstrap replicates; *Stenamoeba* and *Paradermamoeba* served as outgroups. Node values are ML bootstrap percentage / Bayesian posterior probability from a MrBayes analysis of the same alignment under GTR + Γ. Tips are annotated with the substrate and location from which each strain was isolated (more strain information included in Suppl. Data 1). The dotted branch is shortened by a factor of 1/16 (0.0625) for display. *Sappinia lukoli* sp. nov. (SappSF25) is shown in bold.

### 2.2 Aggregative behaviors, eukaryovory, and hijacking of sorocarpic fruiting bodies

As a culture of *S. lukoli* ages, aggregates begin to form at the distal ends of dung fibers, often around the edges of the dung substrate (Fig. 3A-C; Suppl. Video 1). The cells composing the aggregates are typically wrinkled and are shaped like crumpled paper that fit against each other with minimal gaps (Fig. 3D). These aggregates are variable in size and appear more in drying conditions. The cells inside maintain an adhesive quality such that when grazed with an insect needle, a substantial portion of the aggregate adheres to the needle.

**Fig. 3.**
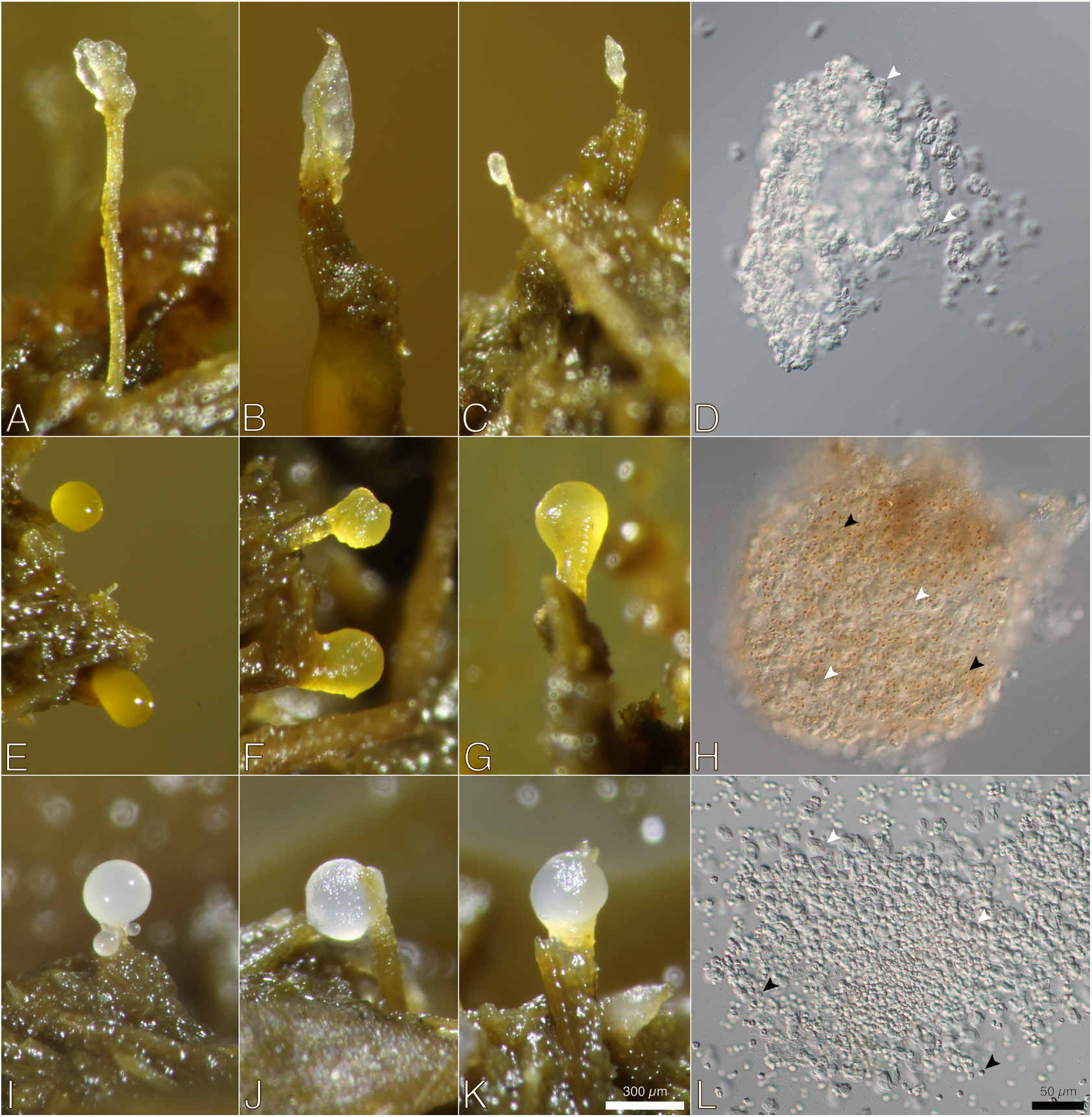
Aggregation and fruiting-body hijacking by *Sappinia lukoli* sp. nov. (A–D) *S. lukoli* cells aggregating at the tips of dung fibers, taken from a co-culture with *Guttulinopsis vulgaris*; aggregated cells are wrinkled and crumpled-paper-like and fit closely together with minimal gaps. (E) Intact sorocarp (fruiting body) of *Sorodiplophrys stercorea*. (F–H) *S. stercorea* sorocarps hijacked by *S. lukoli*. (I) Intact sorocarp of *Guttulinopsis vulgaris*. (J–L) *G. vulgaris* sorocarp hijacked by *S. lukoli*. Hijacking is visible as the sorocarp surface shifts from smooth (E, I) to studded (F, G, J, K), with *S. lukoli* cells distributed throughout the structure rather than only on the surface. In (D, H, L), white arrowheads mark *S. lukoli* cells and black arrowheads mark host spores. Scale bars: white, 300 µm (A, B, C, E, F, G, I, J, K); black, 50 µm (D, H, L).

*S. lukoli* did not grow when offered yeast (*Rhodotorula mucilaginosa*) or bacteria (*Escherichia coli*) on agar, even though we used the same *E. coli* to culture other amoebae including *Sappinia dangeardi* and *Sappinia pedata*. In contrast, *S. lukoli* grew when given the amoebae of *Guttulinopsis vulgaris* or *Sorodiplophrys stercorea* on dung and agar, indicating it is a eukaryovore specialized on other amoebae. However, a wider panel is needed to define its full prey range. Using timelapse microscopy we were able to confirm that these other dung-dwelling species are preyed upon by *S. lukoli* (Suppl. Video 2).

We also consistently find *S. lukoli* cells within the fruiting bodies of these sorocarpic amoebae (Fig. 3E-L). This can be seen by the surface appearance of the fruiting bodies going from smooth (Fig. 3 E, I) to studded (Fig. 3 F, G, J, K). Closer observation of ‘hijacked’ fruiting bodies shows that *S. lukoli* cells are distributed throughout, not just on the surface. In the case of *Sorodiplophrys stercorea* fruiting bodies, most of the *S. lukoli* cells situated inside appeared inactive and showed no visible evidence of eating *S. stercorea* spores. The matrices produced by *S. stercorea* and *G. vulgaris*, along with several of their spores, remained intact, which suggests that *G. vulgaris* spores may be resistant to predation by *S. lukoli* too. Whether *S. lukoli* occupation reduces host spore counts or sorocarp viability remains to be quantified. As the culture continues to age, some *S. lukoli* cells continue to explore across the fruiting bodies and return to the dung substrate, but the majority within seem to keep their position. Internal dynamics within hijacked sorocarps could not be resolved with our current imaging methods and remain to be investigated.

## 3. Discussion

Our results confirm the isolation and cultivation of a molecularly, morphologically, and behaviorally distinct species within the amoebozoan genus *Sappinia*. This adds yet another case of novel protist diversity, and a new predator of sorocarpic amoebae in dung environments. While eukaryovory is broadly established in the sister genus *Thecamoeba* (Dyková et al., 2008; Geisen et al., 2016; Mesentsev and Smirnov, 2021; Page, 1977; Smirnov, 1999), it has not been described in *Sappinia* since Nägler‘s (1909) original account of *S. diploidea* feeding on other amoebae and flagellates. What drives amoeboid species to generalize or specialize on bacterial or eukaryotic prey is not well understood. Considering there are costs associated with generalist diets (Shreenidhi et al., 2024), we suspect this is a worthy area of exploration and the diversity of dung amoebae especially may provide a useful model system.

Historical descriptions of *Sappinia* from dung have noted clusters or heaps of cells forming at the tips of dung fibers, often called a ‘pseudoplasmodium’ (Cienkowski, 1873; Dangeard, 1896; Olive, 1902; Raper, 1960; source text available in Suppl. Data 1). While this is a close behavioral match to *Sappinia lukoli* sp. nov., some records show the constituent cells of the aggregates measure much larger (Cook, 1939). This suggests multiple species of *Sappinia* are capable of forming multicellular aggregates on dung. We expand upon this behavior by recording the capacity of *S. lukoli* to not only aggregate, but to preferentially do so inside the fruiting bodies of sorocarpic amoebae from dung, termed ‘hijacking.’ This novel exploitative interaction signals greater biological complexity and is reminiscent of the dictyostelid *Speleostelium caveatum* (basionym *Dictyostelium caveatum*), which preys on other dictyostelids and interferes with their sorocarpic development (Waddell, 1982; Nizak et al., 2007). However, eukaryote-eukaryote interactions are rarely documented outside the dictyostelids. This represents the first documented case of an aggregative protist colonizing the fruiting bodies of sorocarpic protists across multiple eukaryotic supergroups, all playing out in the same type of habitat, animal dung. That *S. lukoli* cells inside sorocarps largely hold their position, show little movement, and no direct evidence of feeding on host spores raises the possibility that the amoeba is there for occupancy itself, rather than predation. How *S. lukoli* finds host fruiting bodies and preferentially aggregates within remains an open question. They may respond to physical features of the microhabitat, such as the elevated ends of dung fibers that are more dry, where sorocarps are more likely to be. Alternatively, they may follow chemical cues, either from chasing their prey or intercepting the aggregation signals that sorocarpic amoebae secrete to coordinate fruiting body formation. Considering sorocarps are likely arthropod-vectored (Smith et al., 2014), the small cell size and absence of standing morphology in *S. lukoli* may represent trait reductions linked to exploitation of sorocarp-related benefits such as dispersibility. More broadly, recruitment signals are inherently public. The cues used by sorocarpic protists to recruit genetic kin could also be detected and exploited by other lineages, suggesting that cooperation carries an inherent cost (Márquez-Zacarías et al., 2021). *S. lukoli* may represent one of these exploiters, which would make it a tractable system for studying the costs of cooperation among microbes.

Historically, ecological research on dung organisms has focused largely on insects, fungi, and bacteria, with protists receiving comparatively little attention. Yet recent works and metabarcoding surveys show that substantial protist diversity exists in dung from a wide range of animals (Bass et al., 2016). And protists are unlikely to be passive occupants here: in similar terrestrial environments like soils, they act as influential predators that restructure microbial communities and drive morphological and behavioral diversification (Geisen et al., 2018; Matz and Kjelleberg, 2005; Leander, 2020). Here we expand what is known about interactions between dung-dwelling protists, hinting at much unresolved complexity in their ecology and evolution. Beyond the species itself, *Sappinia lukoli* extends what is known of *Sappinia* in several directions: it represents the smallest *Sappinia* described to date, it occupies an undersampled portion of the genus’s SSU rDNA phylogeny where several undescribed strains from soil and soil-adjacent environments cluster, and it reveals another case of aggregative multicellularity. We also reveal further complexity with ecological interactions in the dung landscape in that sorocarps built through coordinated investment by host species can be exploited by phylogenetically distant lineages. Overall, the continued discovery of new species, behaviors, and interactions suggests that herbivore dung, small and ephemeral as it is, is an exciting frontier for protistology.

### Taxonomy of novel species

ZooBank registration number of the present work is urn:lsid: zoobank.org (http://zoobank.org/):pub:XXXXXXXX-XXXX-XXXX-XXXX-XXXXXXXXXXXX

#### *Sappinia lukoli* Henderson & Brown sp. nov

##### Taxonomic summary

Eukaryota (Chatton, 1925) Whittaker & Margulis, 1978

Amorphea Adl et al., 2012

Amoebozoa Lühe, 1913 emend. Cavalier-Smith, 1998

Discosea Cavalier-Smith et al., 2004

Flabellinia Smirnov et al., 2005 *sensu* Kang et al., 2017

Thecavania Jones et al., 2025

Thecamoebida Smirnov and Cavalier-Smith, 2011

Thecamoebidae Schaeffer, 1926

*Sappinia* Dangeard, 1896

*Sappinia lukoli* sp. nov. Henderson & Brown, 2026

##### Diagnosis

*Sappinia lukoli* sp. nov. can be diagnosed by its specific SSU rRNA sequences and by its phylogenetic placement. Additional characters include its relatively small cell size and nuclear dimensions, by the absence of standing behavior under conditions that elicit standing in other *Sappinia* species, and by its aggregation at the tips of dung fibers. Trophozoite cells are lingulate, monopodial, and amoeboid, with lateral folds and occasional dorsal folds during locomotion. On glass, cells moving in a straight line are 18.8–28.2 µm long (average 23.0 µm, n = 75) and 11.4–18.6 µm wide (average 14.6 µm, n = 75), with a length-to-breadth ratio of 1.30–2.05 (average 1.60). The two closely appressed nuclei together span 4.03–5.67 µm at their widest (average 5.14 µm, SD = 0.43, n = 13), and each nucleus is 2.29–3.51 µm in diameter (average 2.86 µm, SD = 0.36, n = 26). Each nucleus usually has a central nucleolus 1.45–2.55 µm in diameter (average 1.93 µm, SD = 0.28, n = 30), and nuclei are rounded rather than hemispherical. Trophozoite cells form multiple small contractile vacuoles, merging into one larger vacuole before expulsion (termed “polyvacuole”) as seen in other *Sappinia* species. Standing cells were not observed under conditions that elicit standing in *S. pedata, S. dangeardi*, and *S. platani*. Aggregative forms consistently occur at the distal tips of dung fibers and, cells can be found within the sorocarps of *Sorodiplophrys stercorea* and *Guttulinopsis vulgaris*. Cysts were not observed. It is primarily a eukaryovore, with documented predation on *S. stercorea* and *G. vulgaris* trophozoites.

##### Type location

Strain SappSF25 of *Sappinia lukoli* sp. nov. was obtained from *Bos taurus* dung at the H. H. Leveck Animal Research Center beef cattle farms at Mississippi State University in Starkville, Mississippi, USA, Lat. 33.4197° N, Long. 88.7953° W, in December of 2025. The strain was originally co-isolated from a *Sorodiplophrys stercorea* sorocarp on the dung surface.

##### Type material

Living cultures of strain SappSF25 are cryopreserved in liquid nitrogen at the laboratory of M. W. Brown, Mississippi State University, and are available upon request.

##### Gene sequence data

The nearly complete SSU-rRNA gene of the type isolate (SappSF25) is 2,525 bp in length and is deposited in GenBank under accession PZ416977.

##### ZooBank ID

[To be added after acceptance]

##### Etymology

The specific epithet lukoli is derived from the Choctaw word meaning “clustered,” “grouped,” or “gathered,” in reference to the aggregative behavior of the cells at dung fiber tips and within host sorocarps. The species is named in recognition of the Choctaw people, on whose ancestral homelands the type strain was isolated.

##### Differential Diagnosis

*S. lukoli* sp. nov. differs from all other known *Sappinia* isolates by its SSU rRNA gene sequences. Morphologically, *S. lukoli* sp. nov. is smaller than all known recorded *Sappinia* species in both cell and nuclear dimensions (Table 1), and exhibits dorsal folds. Behaviorally, *S. lukoli* sp. nov. shows preferential aggregation within and atop the sorocarpic fruiting bodies of *Sorodiplophrys stercorea* and *Guttulinopsis vulgaris*, as well as forming monoclonal aggregates. It has not been observed to stand under conditions that elicit standing in *S. pedata, S. dangeardi*, and *S. platani*, nor have cysts been observed. *S. lukoli* sp. nov. is also confirmed to exhibit eukaryovory, but bacterivory is not excluded.

## 4. Materials and Methods

### 4.1 Sample Collection, Isolation, and Culturing

*Bos taurus* dung was collected from the H. H. Leveck Animal Research Center beef cattle farms at Mississippi State University (coordinates: 33.4197° N, 88.7953° W) and separated into plastic sandwich bags. The dung was then placed into covered glass bowls and incubated at room temperature for 2–3 days. Afterwards, the dung surface was scanned at 5X magnification on a Leica M205C stereoscope. A sorocarp of *Sorodiplophrys stercorea* was carefully picked from the tips of dung fibers using a flame sterilized stainless steel Minutien Insect Pin (Carolina Biological, Burlington, NC, USA), so as to avoid touching the dung below. To sub-culture cells, we placed this sorocarp onto sterilized cow dung bedded onto spring water agar (1 L Ozarka spring water and 15 g agar) with *Escherichia coli* (TOP10 strain MC1061) smeared on top as a prey source. This sterilized dung was prepared by autoclaving 0.5 kg of fresh cow dung in a 1 L beaker covered with aluminum foil. To establish monocultures, *S. stercorea* spores were smeared on the surface of an agar plate, then a single spore was dragged to the edge using a sterilized platinum loop, and this single spore was transferred to fresh culture medium. Likewise, the same was done with *Sappinia lukoli* sp. nov., except we transferred a single cell onto a *S. stercorea* culture, as *S. lukoli* sp. nov. did not grow when given other prey options (*E. coli* and *Rhodotorula mucilaginosa*, a yeast). Cultures were maintained at room temperature, with passages of culture dung with *S. stercorea* and *S. lukoli* sp. nov. cells transferred to fresh media every two weeks.

### 4.2 Microscopy and Morphometrics

To prepare cells for imaging, aggregates of *Sappinia lukoli* sp. nov. and *Sorodiplophrys stercorea* were picked with a flame sterilized insect pin, and placed gently into a drop of OSM (Osmotic Stability Medium) liquid media (2.01 g KCl, 1.58 g NaCl, 0.45 g MgSO_4_ *·*7H_2_O, 0.27 g CaCl_2_ *·*2H_2_O, 1 L deionized water) on a slide and covered with a fresh coverslip. After 5 minutes, the cells had attached to the bottom of the glass slide, and were examined under differential interference contrast (DIC) on a Zeiss Axioskop 2 Plus upright compound microscope (Carl Zeiss Microimaging, Thornwood, NJ, USA). Aggregates were imaged under a 20X Plan-NeoFluar (NA 0.50) and at 2.5X under a Leica M205c stereoscope. Moving cells were imaged under a 40X Plan-NeoFluar (NA 0.75) connected to a Canon (Huntington, NY, USA) CMOS digital camera (EOS R8, 24.2MP full frame mirrorless) controlled by Canon EOS Utility software for Macintosh. Morphometric datasets of *Sappinia lukoli* sp. nov. were curated by selecting for moving cells attached to the glass surface. The pixel measurements of these cells were acquired in ImageJ software (http://imagej.nih.gov/ij/) with the Scale Bar tools for Microscopes utility (http://image.bio.methods.free.fr/ImageJ/?Scale-Bar-Tools-for-Microscopes.html).

### 4.3 Video and Timelapse Microscopy

To record aggregations of *Sappinia lukoli* and their interactions with *Sorodiplophrys stercorea* we used timelapse microscopy. For Supplemental Video 1, we imaged aggregates forming at the edges of culture dung in 3-4 day old cultures using a Canon EOS 650D camera connected to a Leica M205C stereomicroscope taking pictures every 15 seconds. For Supplemental Video 2, to image predation on *S. stercorea*, a small piece of 2-day-old culture dung was placed onto a 0.34 mm thin pad of OSM agar in a glass bottom dish (WillCo Wells, Amsterdam, NL) and agar surface near the edges of the dung was imaged at 20X every 15 seconds. All these image sets were assembled, annotated with timecodes and scale bars, and encoded to MP4 at 30 frames per second using Lapse v1.0.1, open-source software we’ve developed for microscopy datasets (https://github.com/TheBrownLab/Lapse).

### 4.4 Genomic Extraction and SSU rDNA Amplification

Aggregates of *Sappinia lukoli* were picked using a sterilized Minutien insect pin and placed directly into QuickExtract solution (LGC Biosearch Technologies-Epicentre, Madison WI, USA) following the manufacturer’s recommended protocol. This was used as a template for PCR of the full-length 18S rRNA gene with universal eukaryotic SSU primers, 5AmF forward 5’-AACCTGGTTGATCCTGCC (primer S1 in Fiore-Donno et al., 2008) with MedlinB reverse 5’-CCCGGGATCCAAGCTTGATCCTTCTGCAGGTTCAC-CTAC (Medlin et al., 1988) and GoTaq Green Master Mix (Promega). The PCR cycling parameters were 3 m at 95°C, followed by 34 cycles of 30 s at 95°C, 25s at 48°C, and 3.5 min at 72°C. PCR products were purified using Mag-Bind TotalPure NGS magnetic beads (Omega Bio-tek, Inc., Norcross, GA, USA). The purified PCR amplicons were prepared into a library with a Ligation Sequencing DNA V14 Kit (SQK-LSK114) and sequenced on a Nanopore PromethION 2 Solo.

Raw reads were quality- and length-filtered with chopper v0.12.0 (De Coster and Rademakers, 2023), retaining reads with a mean quality *≥*Q10 and a length between 2,200 and 2,800 bp to capture the *Sappinia* 18S while excluding any potential shorter (*∼*1,800 bp) amplified 18S of the co-cultured eukaryote and off-target products. Filtered reads were pooled across PCR replicates and processed with NGSpeciesID v0.3.1 (Sahlin et al., 2021), which clusters reads with isONclust (Sahlin and Medvedev, 2020), generates a per-cluster consensus by racon-polishing (Vaser et al., 2017) a high-quality centroid read, and refines the consensus with medaka v2.2.1 (Oxford Nanopore Technologies, 2018) using the r1041_e82_400bps_sup_v5.2.0 model. The final consensus sequence was verified by BLASTN search (Altschul et al., 1990) against the NCBI nt database. For reference, we deposited SSU rRNA gene sequences for *S. lukoli* (SappSF25), *G. vulgaris* (GvSF24), and *S. stercorea* (SsSF25) strains used in this study in NCBI GenBank (accessions PZ416977, PZ416976, and PZ416975).

### 4.5 Molecular Phylogenetic Analyses

The 18S rRNA gene sequence of *Sappinia lukoli* sp. nov. was analyzed alongside 25 *Sappinia* sequences obtained from GenBank (Clark et al., 2016), with the genera *Stenamoeba* and *Paradermamoeba* included as outgroups (29 total sequences). Locality information for every known *Sappinia* strain is provided as an interactive supplement (Suppl. Data 1), and was used to annotate environmental information on the tree. Sequences were aligned using MAFFT with the L-INS-I algorithm (Katoh and Standley, 2013) and trimmed with BMGE (Criscuolo and Gribaldo, 2010) using a global entropy cutoff (-g) of 0.8. The final alignment was visually inspected for errors and contained 2,368 unambiguously aligned nucleotide characters.

A maximum likelihood tree (Fig. 2) was inferred in RAxML v8.2.12 (Stamatakis, 2014) under the GTRGAMMAI model of nucleotide substitution, with topological support assessed via 1,000 rapid bootstrap replicates. Bayesian inference was performed on the same alignment in MrBayes v3.2.7a (Ronquist et al., 2012) under the GTR + Γ model (nst = 6, rates = gamma). Two independent runs of four chains each were run for 10,000,000 generations, sampling every 500 generations, and the first 25% of samples were discarded as burn-in. Convergence was assessed from the average standard deviation of split frequencies (0.001) and the potential scale reduction factor (PSRF = 1.00 for all parameters). Nodal support in Fig. 2 is given as ML bootstrap percentage / Bayesian posterior probability.

To evaluate sequence divergence within *Sappinia*, we compiled all available *Sappinia* SSU rDNA sequences and realigned them using MAFFT with the AUTO algorithm. This alignment was trimmed with BMGE at a global entropy cutoff (-g) of 0.9 to exclude ambiguously aligned sites. Uncorrected pairwise distances were then calculated in PAUP* (Swofford, 2002) using the “showdist” function (Suppl. Table 1).

## Author contributions

**Tristan C. Henderson:** Writing – review & editing, Writing – original draft, Visualization, Investigation, Formal analysis, Data curation, Conceptualization. **Benjamin Mixon:** Writing – review & editing, Validation, Data curation. **Charles Thompson:** Writing – review & editing, Validation, Data curation. **Cassandra F. van Riessen:** Writing – review & editing, Investigation, Data curation. **Matthew W. Brown:** Writing – review & editing, Investigation, Formal analysis.

## Acknowledgements

This work was supported by the United States National Science Foundation (NSF) Division of Environmental Biology (DEB) grant 2100888 (http://www.nsf.gov) and the Gordon and Betty Moore Foundation (https://doi.org/10.37807/GBMF13832) grant GBMF13832 both awarded to MWB. We thank Mississippi State University for permitting our collection of cattle dung. Lastly, thanks to Virginie M. S. Ruetten for her insightful remarks and thorough reading of this manuscript.

## Data availability

All molecular data and supplemental files (figures and videos) associated with this manuscript, including alignments (trimmed and untrimmed) and phylogenetic trees, will be deposited on FigShare and the DOI added on publication.

## Supplementary Material

**Suppl. Video 1**. Aggregation of *Sappinia lukoli* sp. nov. at the end of a dung fiber. Timelapse of aggregates forming at the edges of culture dung in a 3-4 day old culture.

**Suppl. Video 2**. *Sappinia lukoli* sp. nov. eats *Sorodiplophrys stercorea* cells.

**Suppl. Table 1**. Uncorrected pairwise SSU rDNA distances among all available full-length *Sappinia* SSU sequences.

**Suppl. Data 1**. Interactive supplement giving locality and strain information for all known *Sappinia* strains.

## References

Altschul, S. F., Gish, W., Miller, W., Myers, E. W., and Lipman, D. J. Basic local alignment search tool. Journal of Molecular Biology, 1990. doi: 10.1016/s0022-2836(05)80360-2.

Bass, D., Silberman, J. D., Brown, M., Pearce, R. A., Tice, A. K., Jousset, A., Geisen, S., and Hartikainen, H. Coprophilic amoebae and flagellates, including Guttulinopsis, Rosculus and Helkesimastix, characterise a divergent and diverse rhizarian radiation and contribute to a large diversity of faecal-associated protists. Environmental Microbiology, 18(5):1604–1619, 2016. doi: 10.1111/1462-2920.13235.

Blackwell, M. and Malloch, D. Life History and Arthropod Dispersal of a Coprophilous Stylopage. Mycologia, 83(3): 360, 1991. doi: 10.2307/3759996.

Brefeld, O. Dictyostelium mucoroides. Ein neuer Organismus aus der Verwandtschaft der Myxomyceten. Abhandlungen der Senckenbergischen Naturforschenden Gesellschaft, 7: 85–107, 1869.

Brown, M., Spiegel, F. W., and Silberman, J. D. Phylogeny of the “Forgotten” Cellular Slime Mold, Fonticula alba, Reveals a Key Evolutionary Branch within Opisthokonta. Molecular Biology and Evolution, 26(12):2699–2709, 2009. doi: 10.1093/molbev/msp185.

Brown, M., Silberman, J. D., and Spiegel, F. W. “Slime molds” among the Tubulinea (Amoebozoa): molecular systematics and taxonomy of Copromyxa. Protist, 162(2):277–287, 2011. doi: 10.1016/j.protis.2010.09.003.

Brown, M., Kolisko, M., Silberman, J. D., and Roger, A. J. Aggregative Multicellularity Evolved Independently in the Eukaryotic Supergroup Rhizaria. Current Biology, 22(12): 1123–1127, 2012. doi: 10.1016/j.cub.2012.04.021.

Brown, M. W. and Silberman, J. D. The non-dictyostelid sorocarpic amoebae. In Romeralo, M., Baldauf, S., and Escalante, R., editors, Dictyostelids: Evolution, Genomics and Cell Biology, pages 219–242. Springer, 2013. doi: 10.1007/978-3-642-38487-5_12.

Brown, M. W., Spiegel, F. W., and Silberman, J. D. Amoeba at attention: phylogenetic affinity of Sappinia pedata. Journal of Eukaryotic Microbiology, 54(6):511–519, 2007. doi: 10.1111/j.1550-7408.2007.00292.x.

Cienkowski, L. S. Soobshchenie o neskol’kikh naidennykh im protoplazmaticheskikh organizmakh, predstavlyayushchikh kak by uproshchennyi tip miksomitsetov [Report on several protoplasmic organisms representing a simplified type of myxomycete]. In Levakovskii, N. F., editor, Trudy Chetvertogo S”ezda Russkikh Estestvoispytatelei v Kazani, Vyp.1, Otdelenie Botaniki, Anatomii i Fiziologii Rastenii [Proceedings of the Fourth Congress of Russian Naturalists at Kazan, Part 1, Section of Botany, Anatomy and Plant Physiology], pages 10–11. Kazan, 1873. URL https://search.rsl.ru/ru/record/01003881335.

Clark, K., Karsch-Mizrachi, I., Lipman, D. J., Ostell, J., and Sayers, E. W. GenBank. Nucleic Acids Research, 44(D1): D67–D72, 2016. doi: 10.1093/nar/gkv1276.

Cook, W. R. I. Some observations on Sappinia pedata Dang. Transactions of The British Mycological Society, 22:302–306, 1939. doi: 10.1016/s0007-1536(39)80053-5.

Corsaro, D., Wylezich, C., Walochnik, J., Venditti, D., and Michel, R. Molecular identification of bacterial endosymbionts of Sappinia strains. Parasitology Research, 116:549–564, 2017. doi: 10.1007/s00436-016-5319-4.

Criscuolo, A. and Gribaldo, S. BMGE (Block Mapping and Gathering with Entropy): a new software for selection of phylogenetic informative regions from multiple sequence alignments. BMC Evolutionary Biology, 2010. doi: 10.1186/1471–2148-10-210.

Dangeard, P. Contribution à l’étude des Acrasiées. Le Botaniste, 5:1–20, 1896.

De Coster, W. and Rademakers, R. NanoPack2: population-scale evaluation of long-read sequencing data. Bioinformatics, 39(5):btad311, 2023. doi: 10.1093/bioinformatics/btad311.

Drechsler, C. A Few New Zoöpagaceae Destructive to Large Soil Rhizopods. Mycologia, 31(2):128–153, 1939. doi: 10.1080/00275514.1939.12017328.

Dyková, I., Fiala, I., Dvořáková, H., and Pecková, H. Living together: The marine amoeba Thecamoeba hilla Schaeffer, 1926 and its endosymbiont Labyrinthula sp. European Journal of Protistology, 44(4):308–316, 2008. doi: 10.1016/j.ejop.2008.04.001.

Fiore-Donno, A. M., Meyer, M., Baldauf, S. L., and Pawlowski, J. Evolution of dark-spored Myxomycetes (slime-molds): Molecules versus morphology. Molecular Phylogenetics and Evolution, 2008. doi: 10.1016/j.ympev.2007.12.011.

Fiore-Donno, A. M., Nikolaev, S. I., Nelson, M., Pawlowski, J., Cavalier-Smith, T., and Baldauf, S. L. Deep phylogeny and evolution of slime moulds (Mycetozoa). Protist, 161 (1):55–70, 2010. doi: 10.1016/j.protis.2009.05.002.

Geisen, S., Koller, R., Hünninghaus, M., Dumack, K., Urich, T., and Bonkowski, M. The soil food web revisited: Diverse and widespread mycophagous soil protists. Soil Biology and Biochemistry, 94:10–18, 2016. doi: 10.1016/j.soilbio.2015.11.010.

Geisen, S., Mitchell, E. A. D., Adl, S., Bonkowski, M., Dunthorn, M., Ekelund, F., Fernández, L. D., Jousset, A., Krashevska, V., Singer, D., Spiegel, F. W., Walochnik, J., and Lara, E. Soil protists: a fertile frontier in soil biology research. FEMS Microbiology Reviews, 42(3):293–323, 2018. doi: 10.1093/femsre/fuy006.

Gelman, B. B., Rauf, S. J., Nader, R., Popov, V., Borkowski, J., Chaljub, G., Nauta, H. J. W., and Visvesvara, G. S. Amoebic encephalitis due to Sappinia diploidea. JAMA, 285(19): 2450–2451, 2001. doi: 10.1001/jama.285.19.2450.

Goodfellow, L. P., Belcher, J. H., and Page, F. C. A light- and electron-microscopical study of Sappinia diploidea, a sexual amoeba. Protistologica, 10(2):207–216, 1974.

Gowdham, P. Naegleria fowleri (brain-eating amoeba): a comparative epidemiological and pathophysiological review: global, Indian, and Kerala perspectives. Nightingale, Kauvery Hospital, 2025. URL https://www.kauveryhospital.com/nightingale/naegleria-fowleri-review-2025/.

Henderson, T., Garcia-Gimeno, L., Beasley, C. E., Fry, N. W., Bess, J., and Brown, M. W. High above the rest: Standing behaviors in amoebae of Sappinia and Thecamoeba. European Journal of Protistology, 2024. doi: 10.1016/j.ejop.2024.126082.

Katoh, K. and Standley, D. M. MAFFT Multiple Sequence Alignment Software Version 7: Improvements in Performance and Usability. Molecular Biology and Evolution, 2013. doi: 10.1093/molbev/mst010.

LamŻa, Ł. Diversity of ‘simple’ multicellular eukaryotes: 45 independent cases and six types of multicellularity. Biological Reviews, 98(6):2188–2209, 2023. doi: 10.1111/brv.13001.

Leander, B. S. Predatory protists. Current Biology, 30(10): R510–R516, 2020. doi: 10.1016/j.cub.2020.03.052.

Márquez-Zacarías, P., Conlin, P. L., Tong, K., Pentz, J. T., and Ratcliff, W. C. Why have aggregative multicellular organisms stayed simple? Current Genetics, 67(6):871–876, 2021. doi: 10.1007/s00294-021-01193-0.

Matz, C. and Kjelleberg, S. Off the hook – how bacteria survive protozoan grazing. Trends in Microbiology, 13(7): 302–307, 2005. doi: 10.1016/j.tim.2005.05.009.

Medlin, L., Elwood, H. J., Stickel, S., and Sogin, M. L. The characterization of enzymatically amplified eukaryotic 16S-like rRNA-coding regions. Gene, 1988. doi: 10.1016/0378-1119(88)90066-2.

Mesentsev, Y. and Smirnov, A. Thecamoeba astrologa n. sp. – A new species of the genus Thecamoeba (Amoebozoa, Discosea, Thecamoebida) with an unusually polymorphic nuclear structure. European Journal of Protistology, 81: 125837, 2021. doi: 10.1016/j.ejop.2021.125837.

Michel, R., Wylezich, C., Hauroeder, B., and Smirnov, A. V. Phylogenetic position and notes on the ultrastructure of Sappinia diploidea (Thecamoebidae). Protistology, 4(4): 319–325, 2006.

Mulec, J., Dietersdorfer, E., Üstüntürk-Onan, M., and Walochnik, J. Acanthamoeba and other free-living amoebae in bat guano, an extreme habitat. Parasitology Research, 115(4):1375–1383, 2016. doi: 10.1007/s00436-015-4871-7.

Nägler, K. Entwicklungsgeschichtliche Studien über Amöben. Archiv für Protistenkunde, 15:1–53, 1909.

Nizak, C., Fitzhenry, R. J., and Kessin, R. H.Exploitation of other social amoebae by Dictyostelium caveatum. PLoS ONE, 2(2):e212, 2007. doi: 10.1371/journal.pone.0000212.

Noble, G. A. Coprozoic protozoa from Wyoming mammals. Journal of Protozoology, 5(1):69–74, 1958. doi: 10.1111/j.1550-7408.1958.tb02528.x.

Olive, E. W. Monograph of the Acrasieae. Proceedings of the Boston Society of Natural History, 30:451–510, 1902.

Oxford Nanopore Technologies. medaka. Computer software, 2018. URL https://github.com/nanoporetech/medaka.

Page, F. C. The genus Thecamoeba (Protozoa, Gymnamoebia). Species distinctions, locomotive morphology, and protozoan prey. Journal of Natural History, 11(1):25–63, 1977. doi: 10.1080/00222937700770031.

Qvarnstrom, Y., Silva, A. J., Schuster, F. L., Gelman, B. B., and Visvesvara, G. S. Molecular Confirmation of Sappinia pedata as a Causative Agent of Amoebic Encephalitis. The Journal of Infectious Diseases, 199(8):1139–1142, 2009. doi: 10.1086/597473.

Raper, K. B. Levels of cellular interaction in amoeboid populations. Proceedings of the American Philosophical Society, 104(6):579–604, 1960.

Ronquist, F., Teslenko, M., van der Mark, P., Ayres, D. L., Darling, A., Höhna, S., Larget, B., Liu, L., Suchard, M. A., and Huelsenbeck, J. P. MrBayes 3.2: Efficient Bayesian phylogenetic inference and model choice across a large model space. Systematic Biology, 61(3):539–542, 2012. doi: 10.1093/sysbio/sys029.

Sahlin, K. and Medvedev, P. De novo clustering of long-read transcriptome data using a greedy, quality value-based algorithm. Journal of Computational Biology, 27(4):472–484, 2020. doi: 10.1089/cmb.2019.0299.

Sahlin, K., Lim, M. C. W., and Prost, S. NGSpeciesID: DNA barcode and amplicon consensus generation from long-read sequencing data. Ecology and Evolution, 11(3):1392–1398, 2021. doi: 10.1002/ece3.7146.

Shadwick, L. L., Spiegel, F. W., Shadwick, J. D. L., Brown, M. W., and Silberman, J. D. Eumycetozoa = Amoebozoa? SSUrDNA phylogeny of protosteloid slime molds and its significance for the amoebozoan supergroup. PLoS ONE, 4 (8):e6754, 2009. doi: 10.1371/journal.pone.0006754.

Shreenidhi, P. M., Brock, D. A., McCabe, R. I., Strassmann, J. E., and Queller, D. C. Costs of being a diet generalist for the protist predator Dictyostelium discoideum. Proceedings of the National Academy of Sciences, 121:e2313203121, 2024. doi: 10.1073/pnas.2313203121.

Sladecek, F. X. J., Hrcek, J., Klimes, P., and Konvicka, M. Interplay of succession and seasonality reflects resource utilization in an ephemeral habitat. Acta Oecologica, 46: 17–24, 2013. doi: 10.1016/j.actao.2012.10.012.

Smirnov, A. V. Re-description of Thecamoeba munda Schaeffer 1926 (Gymnamoebia, Thecamoebidae), isolated from the Baltic Sea. European Journal of Protistology, 35 (1):66–69, 1999. doi: 10.1016/S0932-4739(99)80023-6.

Smith, J., Queller, D. C., and Strassmann, J. E. Fruiting bodies of the social amoeba Dictyostelium discoideum increase spore transport by Drosophila. BMC Evolutionary Biology, 14:105, 2014. doi: 10.1186/1471-2148-14-105.

Stamatakis, A. RAxML version 8: a tool for phylogenetic analysis and post-analysis of large phylogenies. Bioinformatics, 2014. doi: 10.1093/bioinformatics/btu033.

Swofford, D. L. PAUP∗: phylogenetic analysis using parsimony (∗and other methods), version 4.0. Sinauer Associates, Sunderland, Massachusetts, 2002.

Tice, A. K., Silberman, J. D., Walthall, A. C., Le, K. N. D., Spiegel, F. W., and Brown, M. Sorodiplophrys stercorea: Another Novel Lineage of Sorocarpic Multicellularity. Journal of Eukaryotic Microbiology, 63(5):623–628, 2016. doi: 10.1111/jeu.12311.

Tong, K., Bozdag, G. O., and Ratcliff, W. C. Selective drivers of simple multicellularity. Current Opinion in Microbiology, 67:102141, 2022. doi: 10.1016/j.mib.2022.102141.

Tyml, T. and Dyková, I. Sappinia sp. (Amoebozoa: Thecamoebida) and Rosculus sp. (SAR: Cercozoa) isolated from king penguin guano collected in the Subantarctic (South Georgia, Salisbury Plain) and their coexistence in culture. Journal of Eukaryotic Microbiology, 65(6):869–879, 2018. doi: 10.1111/jeu.12500.

van Tieghem, P. Sur quelques Myxomycetes a plasmode agrege. Bulletin de la Societe Botanique de France, 27: 317–322, 1880.

Vaser, R., Sovic, I., Nagarajan, N., and Sikic, M. Fast and accurate de novo genome assembly from long uncorrected reads. Genome Research, 27(5):737–746, 2017. doi: 10.1101/gr.214270.116.

Waddell, D. R. A predatory slime mould. Nature, 298(5871): 464–466, 1982. doi: 10.1038/298464a0.

Wylezich, C., Walochnik, J., and Michel, R. High genetic diversity of Sappinia-like strains (Amoebozoa, Thecamoebidae) revealed by SSU rRNA investigations. Parasitology Research, 105(3):869–873, 2009. doi: 10.1007/s00436-009-1482-1.

Wylezich, C., Kudryavtsev, A., Michel, R., Corsaro, D., and Walochnik, J. Electron Microscopical Investigations of a New Species of the Genus Sappinia (Thecamoebidae, Amoebozoa), Sappinia platani sp. nov., Reveal a Dictyosome in this Genus. Acta Protozoologica, 54(1):45–51, 2015. doi: 10.4467/16890027ap.15.004.2191.

